# The microbial derived bile acid lithocholate and its epimers inhibit *Clostridioides difficile* growth and pathogenicity while sparing members of the gut microbiota

**DOI:** 10.1101/2023.06.06.543867

**Authors:** Samantha C Kisthardt, Rajani Thanissery, Colleen M Pike, Matthew H Foley, Casey M Theriot

## Abstract

*C. difficile* infection (CDI) is associated with antibiotic usage, which disrupts the indigenous gut microbiota and causes the loss of microbial derived secondary bile acids that normally provide protection against *C. difficile* colonization. Previous work has shown that the secondary bile acid lithocholate (LCA) and its epimer isolithocholate (iLCA) have potent inhibitory activity against clinically relevant *C. difficile* strains. To further characterize the mechanisms by which LCA and its epimers iLCA and isoallolithocholate (iaLCA) inhibit *C. difficile,* we tested their minimum inhibitory concentration (MIC) against *C. difficile* R20291, and a commensal gut microbiota panel. We also performed a series of experiments to determine the mechanism of action by which LCA and its epimers inhibit *C. difficile* through bacterial killing and effects on toxin expression and activity. Here we show that epimers iLCA and iaLCA strongly inhibit *C. difficile* growth *in vitro* while sparing most commensal Gram-negative gut microbes. We also show that iLCA and iaLCA have bactericidal activity against *C. difficile,* and these epimers cause significant bacterial membrane damage at subinhibitory concentrations. Finally, we observe that iLCA and iaLCA decrease the expression of the large cytotoxin *tcdA* while LCA significantly reduces toxin activity. Although iLCA and iaLCA are both epimers of LCA, they have distinct mechanisms for inhibiting *C. difficile*. LCA epimers, iLCA and iaLCA, represent promising compounds that target *C. difficile* with minimal effects on members of the gut microbiota that are important for colonization resistance.

**Importance:** In the search for a novel therapeutic that targets *C. difficile*, bile acids have become a viable solution. Epimers of bile acids are particularly attractive as they may provide protection against *C. difficile* while leaving the indigenous gut microbiota largely unaltered. This study shows that iLCA and iaLCA specifically are potent inhibitors of *C. difficile*, affecting key virulence factors including growth, toxin expression and activity. As we move toward the use of bile acids as therapeutics, further work will be required to determine how best to deliver these bile acids to a target site within the host intestinal tract.

## Introduction

*Clostridioides difficile* is an anaerobic, Gram-positive, spore-forming bacterium that causes *C. difficile* infection (CDI), which can produce clinical disease ranging from diarrhea and pseudomembranous colitis to death [1]. In the United States, CDI has been designated as an urgent public health threat; in 2017 alone there were approximately 500,000 cases and 20,500 deaths [2]. Antibiotic usage is a key risk factor for CDI, with other factors such as recent hospitalization and advanced age creating further risk [3]. While treatment with antibiotics such as vancomycin, and more recently fidaxomicin can resolve CDI, up to 35% of patients will experience disease recurrence after vancomycin treatment, and 19.7% will experience relapse after fidaxomicin treatment [4–6]. Fecal microbiota transplantation (FMT) is a viable last resort treatment option for patients who have had multiple CDI relapses, however transmission of antimicrobial resistant bacteria has been documented in a few cases post FMT, bringing into question the safety and long-term consequences of a FMT [7, 8].

Antibiotic usage is directly linked to initial and recurring CDI because of its effects on the gut microbiota. The indigenous gut microbiota provides colonization resistance against *C. difficile* through different mechanisms including competition for nutrients, and the production of inhibitory metabolites such as secondary bile acids [9–12]. Within the gut, antibiotics indiscriminately eliminate many microorganisms that play important roles in colonization resistance against *C. difficile.* Antibiotic-induced changes in the gut microbiota and metabolome have been described, and secondary bile acid production is directly impacted [10, 13, 14]. Secondary bile acids are produced by the modification of primary bile acids, which are synthesized in the liver, and normally aid in digestion [15]. The primary bile acids cholic acid (CA) and chenodeoxycholic acid (CDCA) are conjugated with a glycine or taurine at the carboxyl group of carbon 24 [16]. During digestion, these conjugated primary bile acids are released from the gall bladder into the small intestine. While most primary bile acids released are recycled by enterohepatic recirculation in the terminal ileum, a small portion escape recycling and are further modified by gut microbes in the colon. These transformations include deconjugation, dehydroxylation, and epimerization, and a newly discovered function, reconjugation [17–20].

Bile salt hydrolases (BSHs) are encoded by many bacteria in the small and large intestine, and catalyze the hydrolysis of the N-acyl bond at carbon 24 of primary conjugated bile acids, producing a deconjugated bile acid and a free glycine or taurine [16]. BSHs play a key role in bile acid biotransformations as they are the first step in this process or the “gatekeepers” of bile acid modifications [21]. After modification by BSHs, deconjugated primary bile acids can undergo further biotransformations by other gut bacteria. Recently, BSHs have been associated with reconjugation of bile acids with many different amino acids, producing microbial conjugated bile acids (MCBAs) such as Phenylalanine-cholic acid, Tyrosine-cholic acid, and Leucine-cholic acid [17–19, 22]. The presence of BSH enzymes and their products (deconjugated bile acids and or MCBAs) are inhibitory to many bacteria including *C. difficile* [17, 18, 21]. Dehydroxylation is performed by far fewer gut bacteria that contain the bile acid inducible (*bai*) operon and occurs at the 7-alpha or 7-beta carbons. The secondary bile acids deoxycholic acid (DCA) and lithocholic acid (LCA) are both produced by dehydroxylation, and possess increased hydrophobicity and toxicity against *C. difficile* compared to primary bile acids [20, 23]. Further modifications such as epimerization are performed by bacteria containing hydroxysteroid dehydrogenases (HSDH), and produce additional secondary bile acids such as the LCA epimers isoLCA (iLCA) and isoalloLCA (iaLCA) [15, 23, 24].

Microbial-mediated bile acid modifications can have a great impact on the composition of the bile acid pool and colonization resistance against pathogens such as *C. difficile* [14]. Of note, secondary bile acids can affect *C. difficile* at several key stages of its lifecycle, including spore germination and vegetative cell outgrowth [25, 26]. Recent work has also shown that some bile acids can bind the large cytotoxin TcdB, directly neutralizing a key virulence factor [27]. Together, these data support the hypothesis that secondary bile acids are important for the inhibition of key aspects of *C. difficile*’s lifecycle and virulence.

LCA is one of the most abundant secondary bile acids in the gut, having been measured at concentrations up to 700 µM in the human cecum [28–30]. Previous work done by our lab and others has shown that LCA and its epimer iLCA affect aspects of the *C. difficile* life cycle including spore germination, growth, and toxin activity [24, 26]. In particular, epimers of LCA are effective in lowering *C. difficile* carriage and shedding *in vivo*, as well as inhibiting other Gram-positive bacteria in the gut [24]. Bile acid epimers have lower detergent activity and are less toxic to host cells, indicating their potential for use as antimicrobials [24, 31]. LCA epimers iLCA and iaLCA also possess immunomodulatory properties and influence T cell differentiation, suggesting that LCA and its epimers have anti-inflammatory properties that can be leveraged to control inflammation during CDI [32, 33]. With this in mind, we hypothesize that the secondary bile acid epimers iLCA and iaLCA will have stronger inhibitory activity against *C. difficile* than their precursor LCA.

In order to evaluate the therapeutic properties of LCA and its epimers iLCA and iaLCA, we measured their ability to affect each stage of the *C. difficile* lifecycle, and the growth of key members of the indigenous gut microbiota. We show that LCA and its epimers alter different aspects of the *C. difficile* lifecycle including growth, toxin expression, and activity, while sparing mostly Gram-negative members of the gut microbiota. We found that epimers of LCA were potent against *C. difficile* while maintaining host cell integrity. This provides further evidence as bile acids undergo biotransformations by the gut microbiota, especially LCA derivatives, their potency against *C. difficile* increases. Understanding how these new secondary bile acid derivatives affect *C. difficile*, members of the gut microbiota, and the host will be important for designing precision therapeutics for the prevention and treatment of CDI.

## Methods

### *C. difficile* bacterial cultures

*C. difficile* R20291 spores were plated on BHI (brain heart infusion) agar (BD, cat # 211059) with 10 mg/L L-cysteine (MP Biomedical cat # 101444) and 10% weight/volume taurocholate (Sigma Aldrich, cat # T4009) and incubated at 37°C overnight. One colony of germinated spores was inoculated into 10 mL of BHI broth (BD, cat # 101444) with 10 mg/mL L-cysteine and incubated at 37 °C for 16 hr to create a starter culture. After incubation, the starter culture was diluted 1:5 and 1:10 into fresh BHI + cysteine, the optical density (OD_600_) was measured (WPA Biowave CO8000, VWR cat. # 490005-906), and the diluted cultures were allowed to double for 2-3 hr which was confirmed by OD_600_ until the cells reached mid-log. The culture was then diluted to the desired final OD_600_ for experimental use.

### Preparation of bile acids

Bile acid solutions were prepared by dissolving the appropriate concentration of each bile acid in 1 mL of solvent (ethanol or DMSO) to produce a stock solution. This solution was then further diluted in culture to reach the desired final concentration. Bile acids used are as follow (Steraloids catalog ID: TCDCA (C0990-000), GCDCA (C0960-000), CDCA (C0940-000), LCA (C1420-000), 3-oxo LCA (C1750-000), iLCA (C1475-000), iaLCA (C0700-000) (see Table 1).

**Table 1.** Minimum inhibitory concentration for *C. difficile* strain R20291 against bile acids Bile acid MIC (mM)

| Bile acid | MIC (mM) |
| --- | --- |
| Taurochenodeoxycholic acid (TCDCA) | >25 |
| Glycochenodeoxycholic acid (GCDCA) | >6.25 |
| Chenodeoxycholic acid (CDCA) | 1.25 |
| Lithocholic acid (LCA) | 1.5 |
| 3-oxo-lithocholic acid (3-oxo LCA) | 0.1 |
| iso-lithocholic acid (iLCA) | 0.03 |
| iso-allo-lithocholic acid (iaLCA) | 0.02 |

### Minimum inhibitory concentration assays

Minimum inhibitory concentration (MIC) was determined using a modified broth microdilution method as previously described [34]. Culture medium used for all *Clostridia* sp. and *Escherichia coli* was BHI supplemented with 100 mg/L L-cysteine. *Bacteroides* were grown in the same media supplemented with 100 mg/L L-cysteine and 2 µM hemin. *Lactobacillus* sp. were grown in MRS. *Bifidobacterium infantis* was grown in MRS supplemented with 500 mg/L L-cysteine. *M. intestinale* DSM 28989, *P. intestinale* DSM 100749, and *D. muris* DSM 107170 were cultured anaerobically in Fastidious Anaerobe Broth (Neogen) supplemented with 0.01 mg/mL Vitamin K3 (menadione), 0.001 mg/mL Vitamin B12, 5 mg/mL cysteine, and 19 uM hematin. The inoculums were prepared by culturing the isolates in the respective broths at 37°C in an anaerobic growth chamber. After 14 hr of growth, cultures were sub-cultured at a 1:10 dilution and allowed to grow for 3-4 hr anaerobically at 37 °C. The cultures were then diluted in fresh BHI with 100 mg/L L-cysteine so that the starting inoculum OD_600_ was 0.01. Cultures were grown in 96-well plates with each well containing 200 µL of broth supplemented with 2.5% of the varying concentrations of bile acids. Bile acid concentrations ranged from 0.05 mM to 25 mM for TCDCA, 0.05 mM to 6.25 mM for GCDCA, 0.05 mM to 1.5 mM for LCA and 0.01mM to 1.25 mM for iLCA, 3-oxo LCA, and iaLCA. Positive controls included cultures containing no bile acid or containing only solvent (2.5 % DMSO). Uninoculated test media for each strain was used as a negative control to check for sterility. The assay plates were then sealed using a sterile polyester film (VWR, cat # 89134-432) before placing the lid to prevent the panel from dehydrating during incubation. All isolates except *Clostridium scindens*, *Clostridium hiranonis*, and *Clostridium hylemonae* were incubated for 24 hr, whereas the three commensal *Clostridia*, were allowed to grow for 48 hr. MICs were defined as the lowest concentration of the bile acid supplement at which there was no growth as measured by optical density. The titer of the initial inoculum was determined by plating on agar (BHI with 100 mg/L L-cysteine for *Clostridia* and *E. coli,* and BHI with 100 mg/L L-cysteine and 2 µM hemin for *Bacteroides*. MRS plates for *Lactobacillus* sp., and MRS supplemented with 500 mg/L L-cysteine for *B. infantis*). After 24 hr of incubation the same dilutions with countable colonies as identified from the initial inoculum were plated to determine growth inhibition.

### Time dependent kill assay

Time dependent kill assay of *C. difficile* was performed using a modified kill kinetics assay protocol as described previously [34]. Briefly, overnight *C. difficile* cultures were back-diluted 1:10 into pre-reduced BHI plus 100 mg/L L-cysteine broth and allowed to grow until OD_600_ reached mid log (0.45–0.50). The *C. difficile* cells were challenged with or without bile acids (1X and 5X MIC of iLCA and iaLCA, and 1X MIC of LCA) or DMSO (solvent). All tubes were incubated anaerobically at 37°C for 24 hr. At intervals of 0, 2, 4, and 24 hr, aliquots were removed and serially diluted. Unheated samples were plated on either BHI agar containing 100 mg/L L-cysteine to enumerate only vegetative cells, or on TBHI agar (BHI containing 100 mg/L L-cysteine and 0.1% taurocholate) to enumerate total vegetative cells and spores. After heat treatment (65 °C for 20 min) samples were plated on TBHI to enumerate total spores. Colonies were counted the next day to determine colony forming units per mL (CFU/mL). Experiments were performed with 3 biological replicates.

### Membrane integrity assay

Exponentially growing *C. difficile* was captured mid log (OD_600_ = 0.8) during growth in BHIS with 100 mg/L L-cysteine, then back-diluted to a final OD_600_= 0.1 into BHIS containing a given bile acid for 3 hr. Bile acid supplements (5 µL) at 0.25X and 0.5X MIC of LCA, iLCA, and iaLCA were added to 195 µL of culture. Tubes containing *C. difficile* cultures with no supplement or heat killed *C. difficile* cultures were included as controls. Following exposure, cultures were stained with propidium iodine (PI) using slight modifications to a previous method [31, 35]. Bacteria were heat-killed at 80°C for 10 min as positive controls for PI staining. PI fluorescence was measured from flat-bottom clear plates (excitation: 540 nm; emission: 610 nm) and was normalized to OD_600_.

### *C. difficile* toxin expression assay

For toxin expression assays, *C. difficile* R20291 carrying either pDSW1728-P*tcdA*::*mCherry* or pDSW1728 was plated on BHI containing 10 mg/L L-cysteine and 10 µg/mL thiamphenicol (Sigma, cat # T0261). After an overnight incubation at 37°C, one colony was inoculated into 10 mL of BHI broth with 10 mg/L L-cysteine and 10 µg/mL thiamphenicol and incubated at 37°C for 14 hr. This starter culture was diluted 1:5 and 1:10 into fresh BHI + cysteine + thiamphenicol and allowed to double for approximately 4 hr. Experimental cultures were diluted to a starting OD_600_ of 0.01. After incubation, a modified protocol based on Ransom et al. [36] was used to fix the cultures for fluorescence measurement. Briefly, a 5x fixation master mix was prepared fresh in aerobic conditions and aliquoted into 1.5 mL Eppendorf tubes. These tubes were passed into the anaerobic chamber (Coy, USA) and 500 µL of culture was added to each tube containing fixative. The tubes containing culture were removed from the chamber, and incubated and fixed as directed by Ransom et al [36]. After fixation, fluorescence was measured following the protocol as described by Ransom et al. A Tecan plate reader (Tecan, Switzerland) was used along with Magellan software (Magellan, V7.2) to obtain fluorescence and absorbance measurements [36]. A black 96 well plate (Genesee Scientific cat # 91-424TB) was used to measure the fluorescence and a clear 96 well plate (Genesee Scientific, cat # 25-104). was used to measure optical density. Relative fluorescence was reported as a ratio of fluorescence to absorbance and graphed with Prism version 9.4.1 (GraphPad Software, LaJolla, CA).

### Vero cell cytotoxicity assay

*C. difficile* was grown overnight at 37°C in pre-reduced BHI plus 100 mg/L L-cysteine broth in an anaerobic chamber. Overnight *C. difficile* cultures were sub-cultured 1:10 into same media, and allowed to grow for 3 hr anaerobically at 37°C. The culture was then diluted in fresh BHI so that the starting OD_600_ was 0.01. To this test compounds LCA, iLCA, and iaLCA were added at 0.25X and 0.5X MIC to make a total volume of 5 mL. Controls included *C. difficile* only cultures and DMSO treated cultures to make a final concentration of 2.5%. All treatments were incubated for 48 hr at 37°C in an anaerobic chamber. At 24 hr, aliquots were removed from each treatment, serially diluted 10-fold in phosphate buffered saline (PBS) and plated on BHI plus 100 mg/L L-cysteine and 0.1% taurocholate using a track dilution method[37]. This method involved plating 10 μL of six dilutions on separate tracks of a single square plate (Genesee Scientific, Cat # 26-275). All plates were incubated at 37°C for 24 hr anaerobically. Plates were counted the next day to enumerate total vegetative cells plus spores in each treatment. At 48 hr all treatments were removed from the chamber and stored in −80°C until use for measuring toxin activity from the culture supernatants.

Toxin activity was measured by a Vero cell cytotoxicity assay [26]. Vero cells were grown and maintained in DMEM media (Gibco Laboratories, 11965-092) with 10% fetal bovine serum (Gibco Laboratories, 16140-071) and 1% Penicillin streptomycin solution (Gibco Laboratories, 15070-063). Cells were incubated with 0.25% trypsin (Gibco Laboratories, 25200-056), washed with 1X DMEM media, and harvested by centrifugation 1,000 RPM for 5 min. Cells were plated at 1 × 10^4^ cells per well in a 96-well flat bottom microtiter plate (Corning, 3596) and incubated overnight at 37°C/5% CO_2_. Growth culture tubes were defrosted on ice and then centrifuged at 1,750 RPM for 5 min to pellet vegetative *C. difficile*. Culture supernatants were collected from each well and serially diluted by 10-fold to a maximum of 10^−6^ using 1X PBS. Sample dilutions were incubated 1:1 with PBS (for all dilutions) or antitoxin (performed for 10^−1^ and 10^−4^ dilutions only, TechLabs, T5000) for 40 min at room temperature. Following incubation, these admixtures were added to the Vero cells. After an overnight incubation at 37°C/5% CO_2_, plates were viewed under 200× magnification for Vero cell rounding. The cytotoxic titer was defined as the reciprocal of the highest dilution that produced rounding in 80% of Vero cells for each sample. Vero cells treated with purified *C. difficile* toxins (A and B) and antitoxin (List Biological Labs, 152C and 155C; TechLabs, T5000) were used as controls. A test cytotoxicity assay was run prior to assays to ensure that the LCA, iLCA, and iaLCA and DMSO did not affect the cytoskeleton of Vero cells at the tested concentrations.

### Cell lines and reagents

Caco-2 cells (ATCC, HTB-37) were cultured in DMEM supplemented with 2 mM L-glutamine and 10% FBS and incubated in 5% CO2 at 37 °C. CellEvent™ Caspase-3/7 Green Detection Reagent (C10723), PrestoBlue™ Cell Viability Reagent (A13261) and Invitrogen™ NucGreen™ Dead 488 ReadyProbes™ Reagent (R37109) were purchased from ThermoFisher Scientific.

### Caspase-3/7 activation and viability assay

Activation of caspases 3 and 7 was assessed using CellEvent Caspase-3/7 Green Detection Reagent according to the manufacturer’s instructions. Caco-2 cells were seeded in 96 well plates for 7 days or until confluent before the addition of fresh DMEM containing bile acids or MeOH. After 24 hr of incubation, the media was replaced with 1X PBS containing 5% FBS and 2 µM of CellEvent™ Caspase-3/7 Green Detection Reagent. The plates were incubated for 1 hr at 37 °C and green fluorescence was detected at excitation/emission wavelengths of 485/530 nm using a ThermoFisher Fluoroskan Plate Reader. PrestoBlue™ Cell Viability Reagent was then added to the wells and incubated for 30 min at 37 °C. Red fluorescence was detected at excitation/emission wavelengths of 550/610 nm.

### Statistical analysis

Statistical testing was performed using Prism 9 for macOS (GraphPad Software, LaJolla, CA). Statistical significance across all conditions of kill kinetics, toxin expression, and cell viability was determined with a two-way ANOVA with Dunnett’s multiple comparisons test to compare to control conditions. Significance of membrane integrity and toxin activity was determined by an unpaired t-test with Welch’s correction. All tests were run with a set statistical significance of p < 0.05 and at least three biological replicates.

## Results

### Microbial epimerization of LCA enhances antibacterial activity against *C. difficile*

As bile acids are modified, they are associated with reduced *C. difficile* growth [9, 14]. To understand how the modification of LCA affects *C. difficile* growth, we used a MIC assay. Using a modified broth microdilution method previously described in Thanissery et al. [34], we measured the MIC of the LCA precursors TCDCA, GCDCA, and CDCA as well as the intermediate 3-oxoLCA, and finally the epimers iLCA and iaLCA against *C. difficile* strain R20291 (Fig. 1A). We observed that the primary conjugated bile acids TCDCA and GCDCA have high MICs of greater than 25 mM and greater than 6.25 mM, respectively. In comparison, the MIC of LCA was 1.5 mM and the MIC of 3-oxo-LCA is 0.1 mM, while iLCA had a MIC of 0.03 mM and iaLCA had a MIC of 0.02 mM (Fig. 1B, Table 1). This indicates that as bile acids undergo biotransformations by the gut microbiota, especially LCA derivatives, their inhibition against *C. difficile* increases.

**Figure 1 –.**
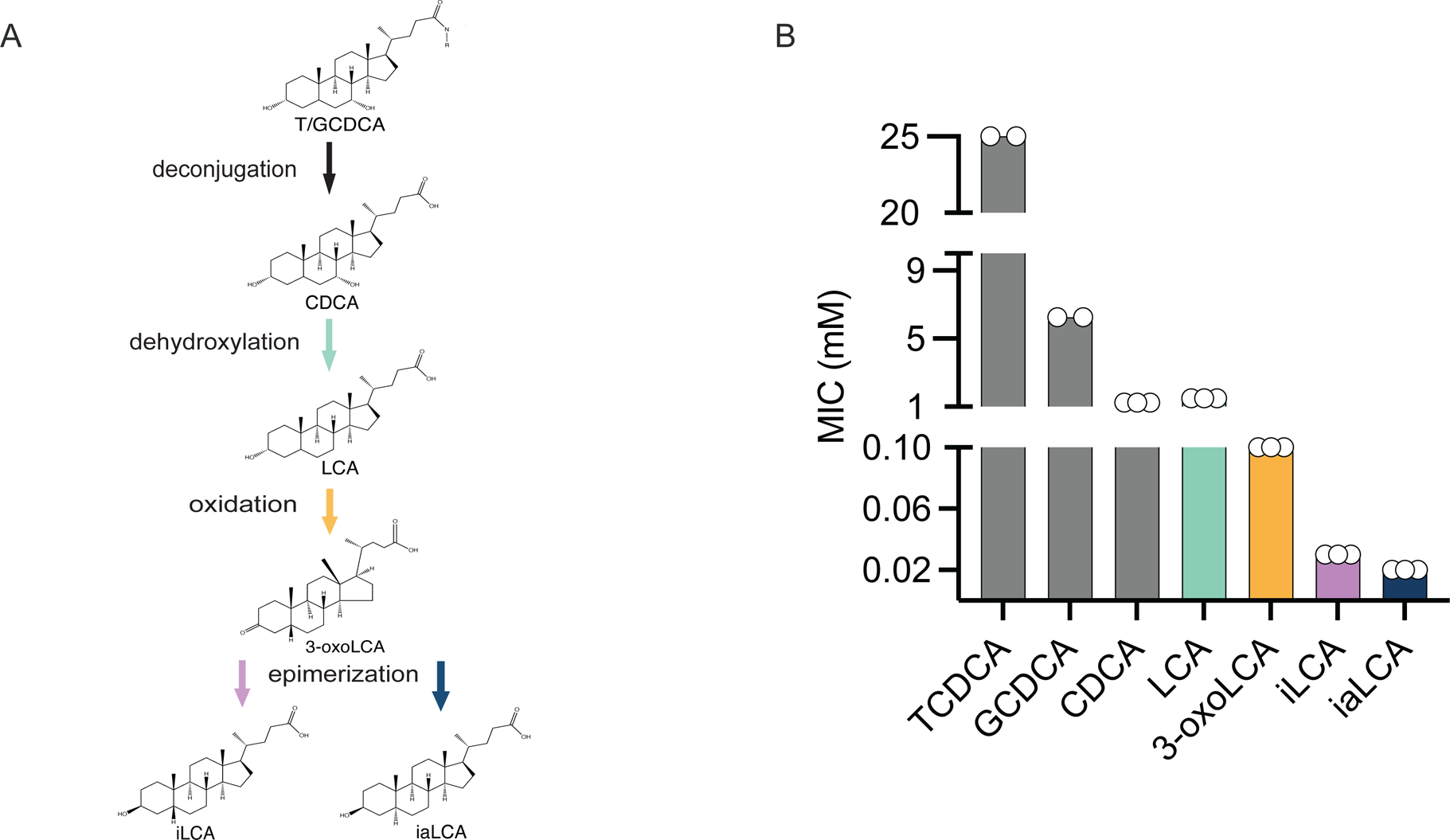
LCA and its derivatives are able to inhibit *C. difficile* growth at very low concentrations. (A) Structures and abbreviations are as follows: taurochenodeoxycholic acid (TCDCA), glycochenodeoxycholic acid (GCDCA), chenodeoxycholic acid (CDCA), lithocholic acid (LCA), 3-oxo-lithocholic acid (3-oxo LCA), iso-lithocholic acid (iLCA), iso-allo-lithocholic acid (iaLCA). (B) Minimum inhibitory concentration (MIC) of all bile acids were found using a modified broth microdilution method. Bile acid modifications by the gut microbiota increase their potency against *C. difficile* R20291. Data presented represents mean ± SEM of n=2-3 samples.

### LCA derivatives inhibit *C. difficile* while sparing members of the indigenous gut microbiota

Ensuring that compounds targeting *C. difficile* do not disrupt other members of the indigenous gut microbiota is essential to reducing the likelihood of recurring disease [38]. Having determined that LCA, iLCA, and iaLCA can inhibit the pathogen *C. difficile* at low concentrations, we wanted to examine how they affected several bacteria commonly found in the gut. We selected twelve bacterial isolates that are considered keystone species and highly abundant in the indigenous gut microbiota, where colonization resistance is intact [34] (Table 2). We additionally selected *C. scindens, C. hylemonae,* and *C. hiranonis* because they are also producers of secondary bile acids including LCA and its epimer isoLCA by means of 7a-dehydroxylation and epimerization [16, 39]. To determine whether LCA and its epimers affect this twelve-member bacterial panel, we repeated our MIC assays with LCA, 3-oxoLCA, iLCA, and iaLCA. We found that these bacteria broadly tolerate LCA and its derivatives. All *Lactobacillus, Bacteroides, Escherichia, Bifidobacterium* species, and *Muribaculaceae* family members tolerated LCA and its epimers with MICs greater than those of *C. difficile* (Table 2). In comparison, non-pathogenic *Clostridia* (*C. scindens, C. hylemonae, C. hiranonis*) were inhibited by some bile acids, as evident by low MICs ranging from 0.1 to 0.02 mM (Table 2). The differences in MICs observed indicates that LCA and its epimers are well-tolerated by members of our twelve-member bacterial panel, especially the Gram-negative bacteria, while the *Clostridia* are more susceptible to these bile acids.

**Table 2.**
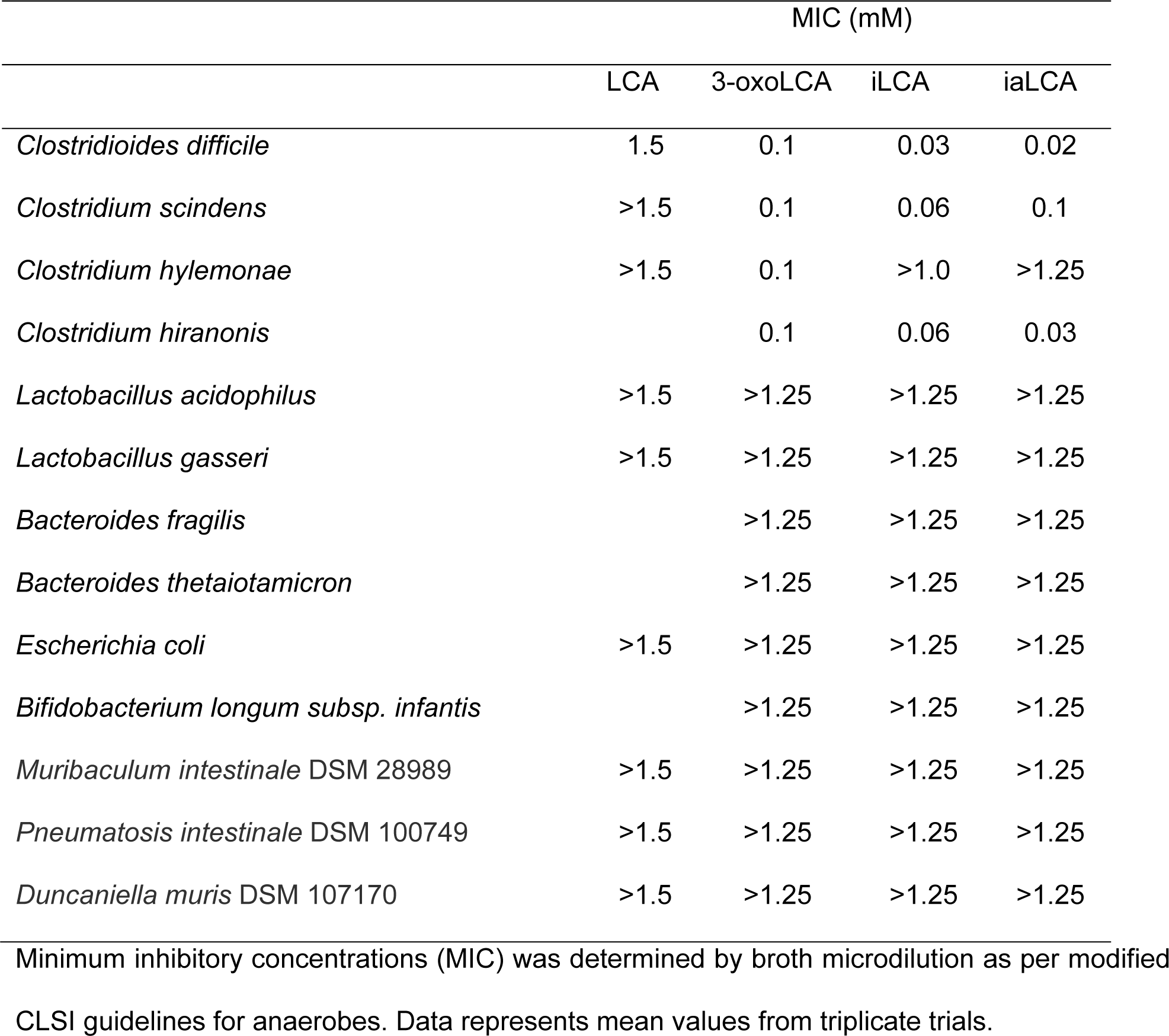
Minimum inhibitory concentration for the commensal panel against LCA and its derivatives

### LCA derivatives cause differential killing of *C. difficile* over a 24 hour period

After determining that LCA and its epimers at low concentrations were able to prevent *C. difficile* growth, we shifted to interrogating how each bile acid affects established growth over time [40]. We continued our experiments with LCA and its epimers iLCA and iaLCA solely because we wanted to address how epimers differ from secondary bile acids in their functionality. Similarly, we did not move forward using 3-oxo-LCA because it is an intermediate form. Using a previously described kill kinetics assay, we asked if LCA, iLCA, and iaLCA could decrease bacterial growth early and late over a 24 hr period [26]. At the MIC, 1.5 mM, only LCA significantly decreased growth, however this was only at the 2- and 4-hr timepoints (LCA compared to untreated control (DMSO) by two-way ANOVA, p <0001, p < 0.001). By 24 hr, growth recovered (Fig. 2A). In comparison, iLCA and iaLCA had no significant effects on killing at the MIC (Fig. 2B, C).

**Figure 2 –.**
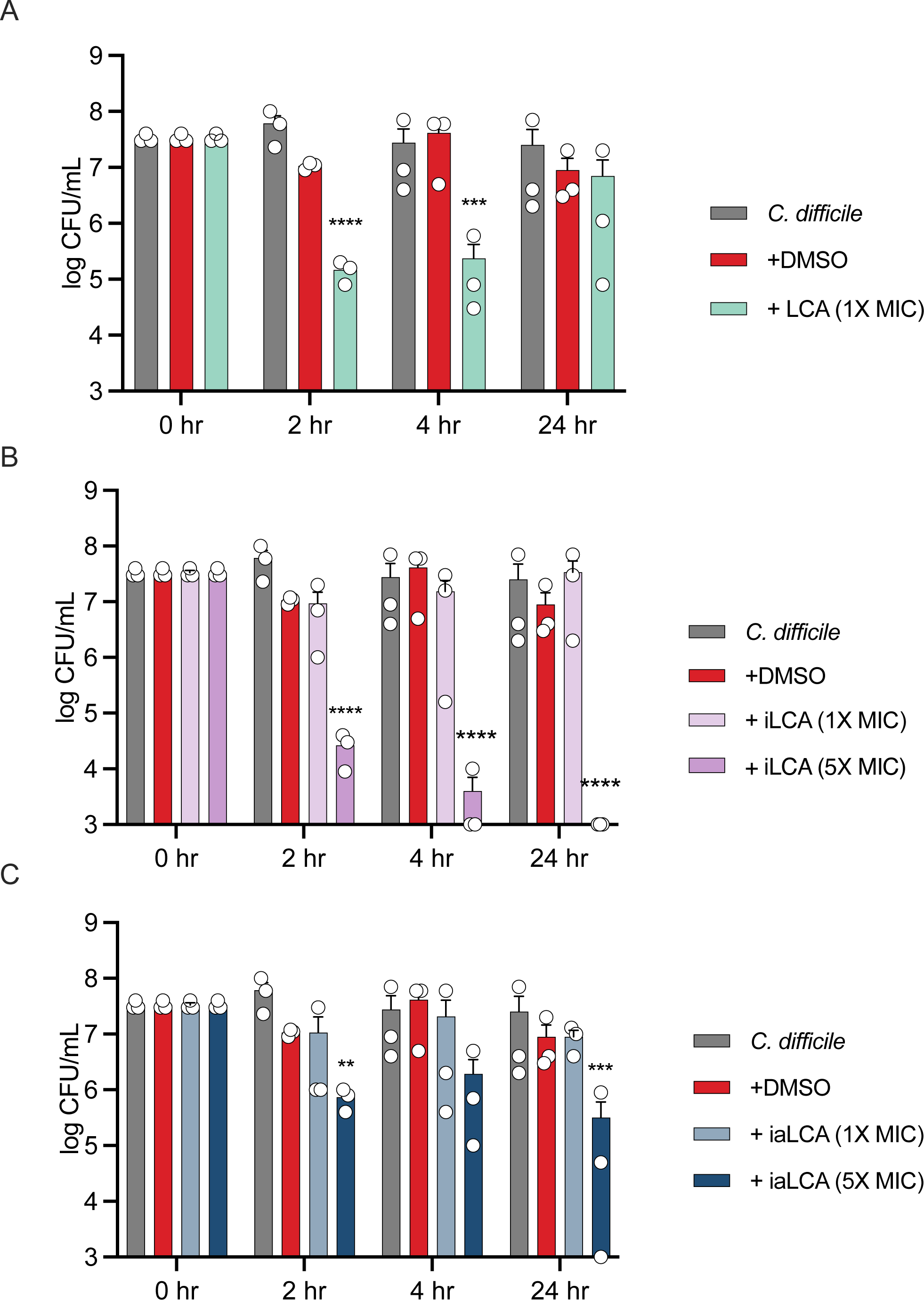
LCA and its derivatives cause differential killing over a 24 hr period. A bacterial kill kinetics assay modified from previous work was used to show killing activity of LCA and its derivatives over a 24 hr period. Effects of (A) LCA and (B) iLCA, and (C) iaLCA on *C. difficile* growth was measured over 24 hr. Statistical significance between DMSO treated control and treatment groups was determined by two-way ANOVA (*, p < 0.05; **, p < 0.01; ***, p < 0.001; ****, p < 0.0001). Data presented represents mean ± SEM of triplicate samples.

To better understand the inhibitory activity of iLCA and iaLCA, we chose to test both bile acids at five times the MIC. Under these conditions, we saw that iLCA caused a consistent and significant decrease in *C. difficile* bacterial load as early as 2 hr (iLCA compared to untreated control (DMSO) by two-way ANOVA, p < 0.0001). In comparison, iaLCA caused a significant decrease in growth at 2 and 24 hr (iaLCA compared to untreated control (DMSO) by two-way ANOVA, p< 0.01, p < 0.001). Taken together, these results suggest that each LCA epimer inhibits *C. difficile* through different mechanisms of action.

### LCA derivatives differentially affect the bacterial membrane permeability of *C. difficile*

Building on the hypothesis that LCA, iLCA and iaLCA may each affect *C. difficile* differently, we began to define how each bile acid alters different parts of the *C. difficile* lifecycle. Since previous work has shown that the detergent-like nature of LCA causes damage to the membrane of some gut bacteria, we sought to define the effects of LCA and its epimers on *C. difficile*’s membrane [41]. Using a propidium iodide-based bacterial membrane permeability assay, we tested the effects of LCA, iLCA, and iaLCA at 0.25X and 0.5X MIC on *C. difficile*. Our controls included a heat-killed sample of *C. difficile,* which represented maximum membrane damage, and two non-heat-killed samples that represent an in-tact cell membrane. One non-heat-killed sample was additionally treated with DMSO, the vehicle in which all bile acids were suspended in. At subinhibitory concentrations, we broadly observed that LCA and its epimers produce differential effects on *C. difficile*’s membrane. When exposed to LCA at 0.25X and 0.5X the MIC, a significant increase in fluorescence relating to membrane damage was seen compared to DMSO-treated controls (LCA compared to untreated control (DMSO) by unpaired t-test with Welch’s correction, p <0.01) (Fig. 3). iLCA did not cause significant membrane damage at 0.25X or 0.5X the MIC. iaLCA, on the other hand, showed a dose dependent response such that it did not cause membrane damage at 0.25X the MIC, but it did at 0.5X the MIC (p < 0.01). These results align with previous work where secondary bile acid DCA has membrane-damaging effects on bacteria and expand on the current paradigm to explore LCA and its epimers iLCA and iaLCA [15, 31].

**Figure 3 –.**
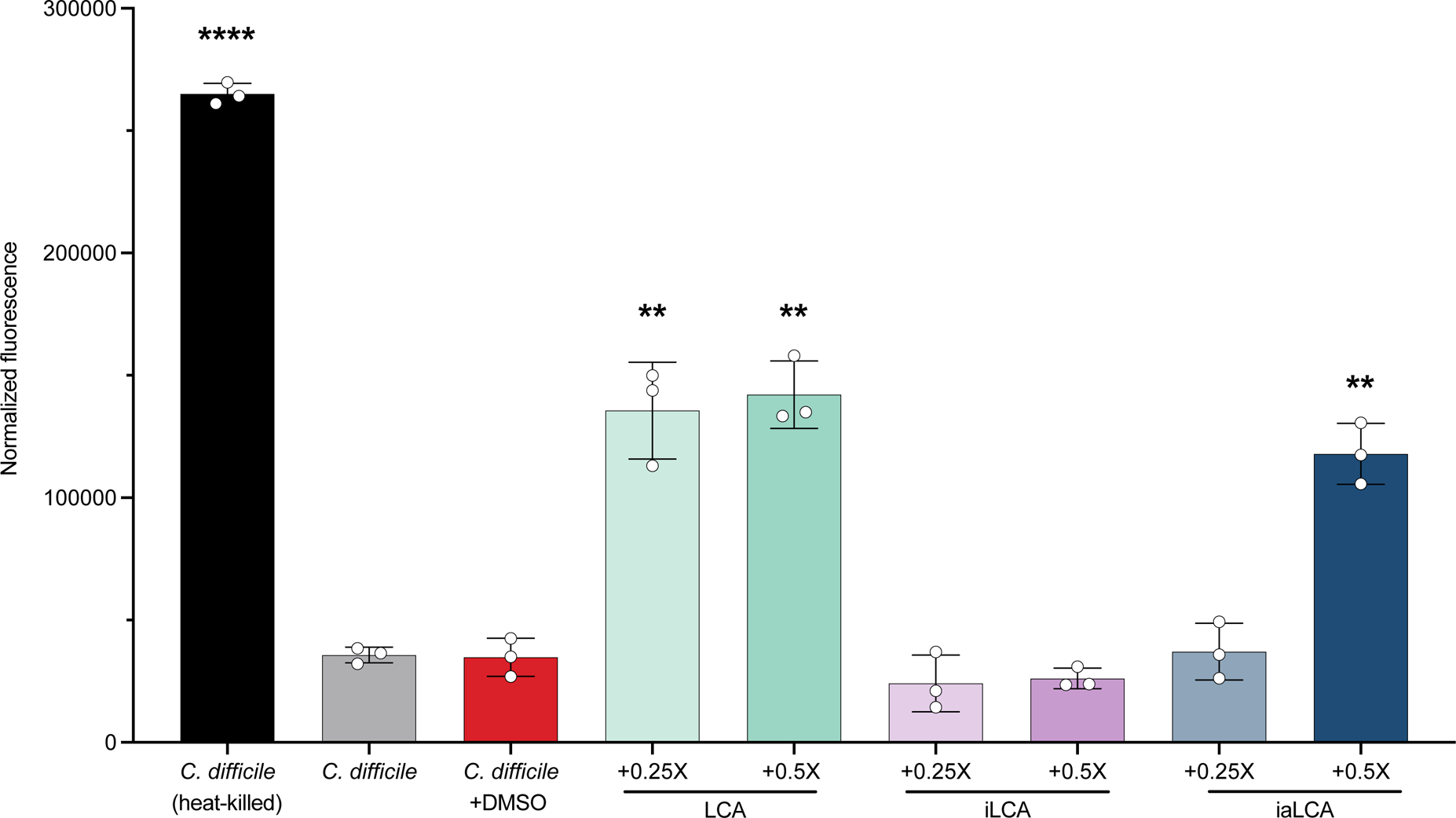
LCA and its derivatives produce differential effects on *C. difficile*’s cell membrane. The effects of LCA, iLCA, and iaLCA at two subinhibitory concentrations were measured on *C. difficile* by a fluorescence-based bacterial membrane viability assay. Statistical significance was determined by unpaired t-test with Welch’s correction (**, p < 0.01; ****, p < 0.0001).

### The LCA derivatives iLCA and iaLCA decrease toxin expression at subinhibitory concentrations

To continue interrogating the potential for differential mechanisms of action behind the inhibition of iLCA and iaLCA, we tested their effects on *C. difficile* toxin expression. The large cytotoxins TcdA and TcdB are significant virulence factors associated with CDI, and previous work has shown that LCA and iLCA have effects on toxin activity [26]. *C. difficile* containing a plasmid construct with the fluorescent reporter mCherry downstream from the *tcdA* promoter was grown in the presence of subinhibitory concentrations of iLCA and iaLCA. LCA was not included as it was found to emit autofluorescence alone and was not compatible with this assay. A control consisting of *C. difficile* containing an empty vector was also grown in these conditions as a negative control. As concentrations of iLCA and iaLCA increased from 0.125X to 0.5X the MIC, expression of *tcdA* significantly decreased (iLCA or iaLCA compared to untreated control (DMSO) by two-way ANOVA, p < 0.001, p < 0.0001) (Fig. 4). This was represented by a decrease in fluorescence in the cultures containing the mCherry reporter. In comparison, no fluorescence was detected in samples containing the empty vector. Fluorescence was reported relative to growth (OD_600_) as a means of normalization. This data indicates that iLCA and iaLCA do influence toxin (*tcdA*) expression.

**Figure 4 –.**
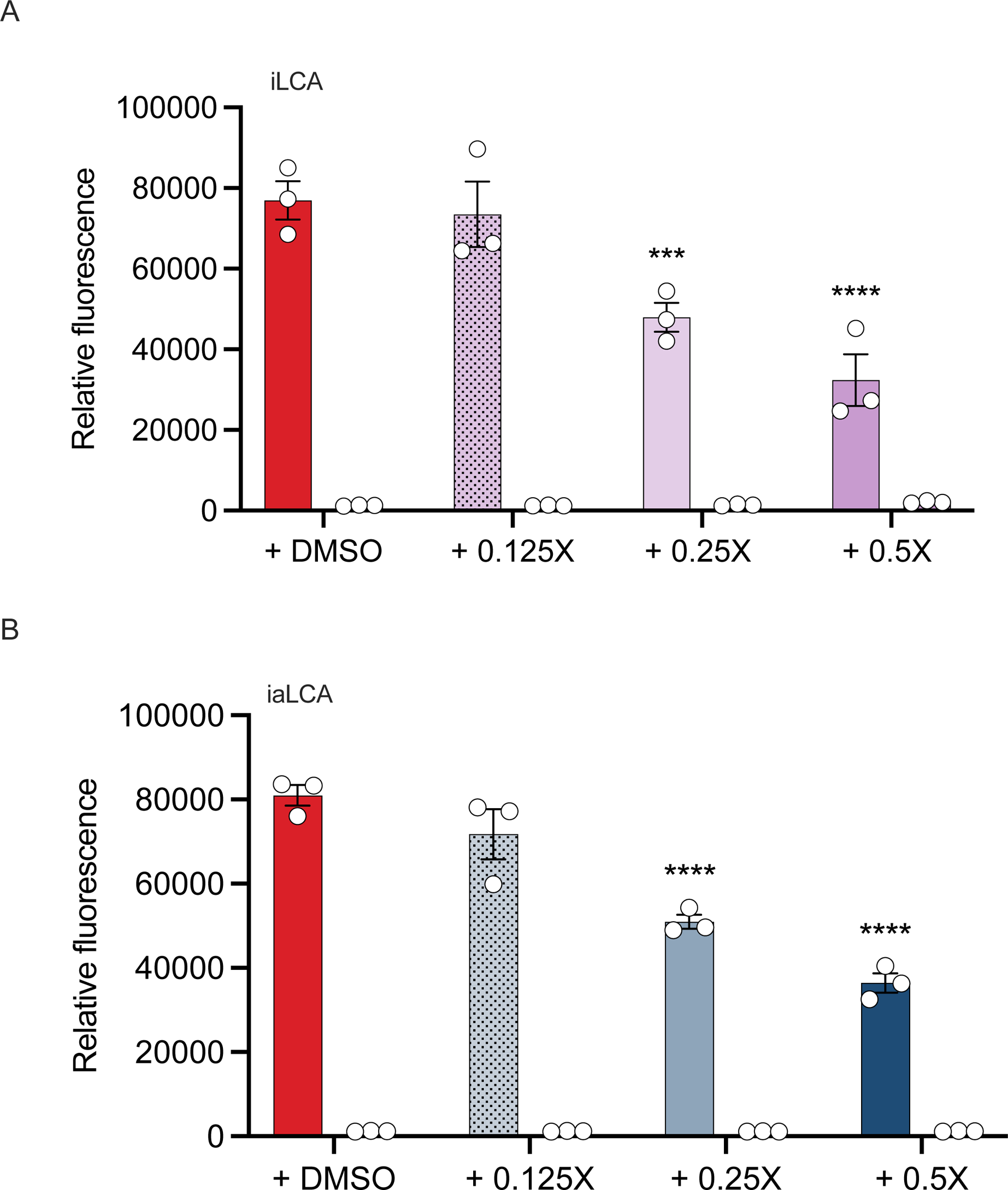
The LCA derivatives iLCA and iaLCA decrease toxin expression at subinhibitory concentrations. Using a reporter strain of *C. difficile* R20291 containing the fluorescent reporter mCherry at the toxin gene *tcdA*, cultures were grown in increasing concentrations of (A) iLCA and (B) iaLCA. An empty vector control was also grown in the same bile acid conditions. Statistical significance between untreated control and treatment groups was determined by two-way ANOVA (***, p < 0.001; ****, p < 0.0001). Data presented represents mean ± SEM of triplicate samples.

### LCA alone decreases toxin activity at subinhibitory concentrations

After observing a decrease in toxin expression when *C. difficile* cultures are supplemented with iLCA and iaLCA, we wanted to know whether toxin activity is also affected. LCA and iLCA have been shown to decrease toxin activity in *C. difficile* strain R20291, leading us to hypothesize that its epimers may behave similarly [26, 27]. To determine if LCA, iLCA, and iaLCA affect toxin activity, we performed a Vero cell cytotoxicity assay. Briefly, *C. difficile* was grown for 48 hr in the presence of 0.25X or 0.5X the MIC of LCA, iLCA, and iaLCA. *C. difficile* grown alone and in the presence of the DMSO vehicle were used as controls. Growth was assessed by bacterial enumeration at 48 hr to determine whether the presence of the bile acids affected bacterial growth as this would also affect toxin activity. Supernatants were then used in a Vero cell cytotoxicity assay.

Subinhibitory concentrations of LCA, iLCA, and iaLCA did not affect *C. difficile* growth at 48 hr (Fig. 5A). Consistent with previous findings, we observed a significant decrease in toxin activity when LCA is supplemented at 0.25X and 0.5X the MIC (Fig. 5B LCA, iLCA, or iaLCA compared to untreated (DMSO) control by unpaired t-test with Welch’s correction, p < 0.05). In comparison, neither concentration of iLCA caused a significant change in toxin activity in *C. difficile.* While iaLCA did not cause a significant change in *C. difficile* toxin activity, it did cause an observable decrease at 0.5X the MIC.

**Figure 5 –.**
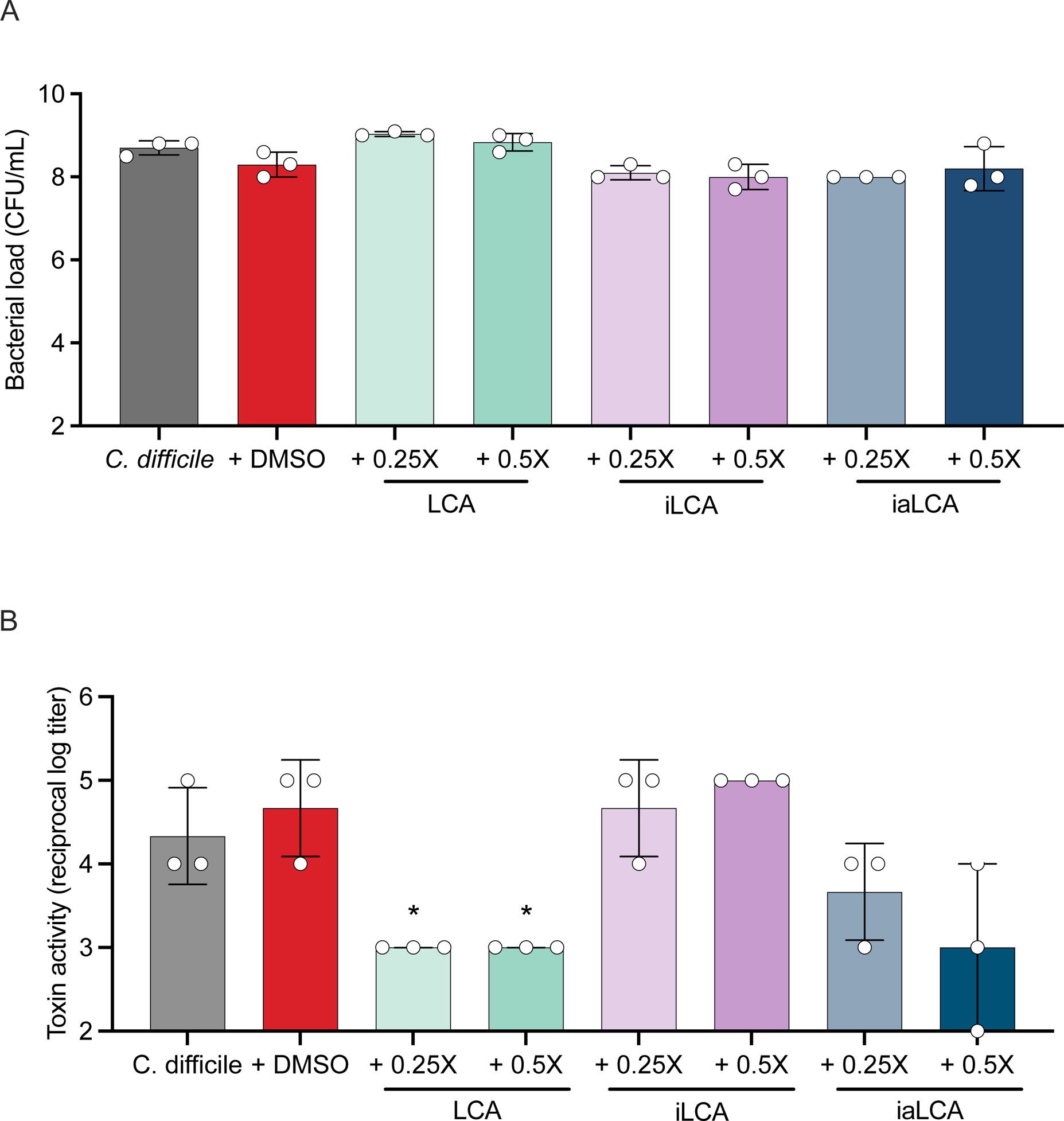
Subinhibitory concentrations of LCA alone decreases *C. difficile* toxin activity while iLCA and iaLCA have differential effects. After 48 hr of growth seen in (A), a Vero cell viability assay (B) was used to assess how the presence of subinhibitory concentrations affect toxin activity in *C. difficile.* Statistical significance between DMSO treated controls and treatment groups was determined by an unpaired t test with Welch’s correction (*, p < 0.05). Data presented represents mean ± SD of samples in triplicate.

### Host cell viability is not affected by LCA epimers

Several bile acids elicit cytotoxic effects against the intestinal epithelium and induce apoptosis and cell death [42]. To test whether LCA and its derivatives could be cytotoxic against host cells, we assessed cell viability and caspase 3/7 activation in Caco-2 cells treated with 0.5X, 1X or 2X MIC of LCA, iLCA and iaLCA. Neither LCA nor its derivatives altered cell viability at 0.5X, 1X and 2X MIC (Fig. 6). LCA induced caspase 3/7 activation at 2X MIC whereas iLCA and iaLCA did not alter apoptosis at any tested MIC (Fig. 6, LCA, iLCA, or iaLCA compared to untreated control (DMSO) by two-way ANOVA **, p < 0.01; ****, p < 0.0001).

**Figure 6 –.**
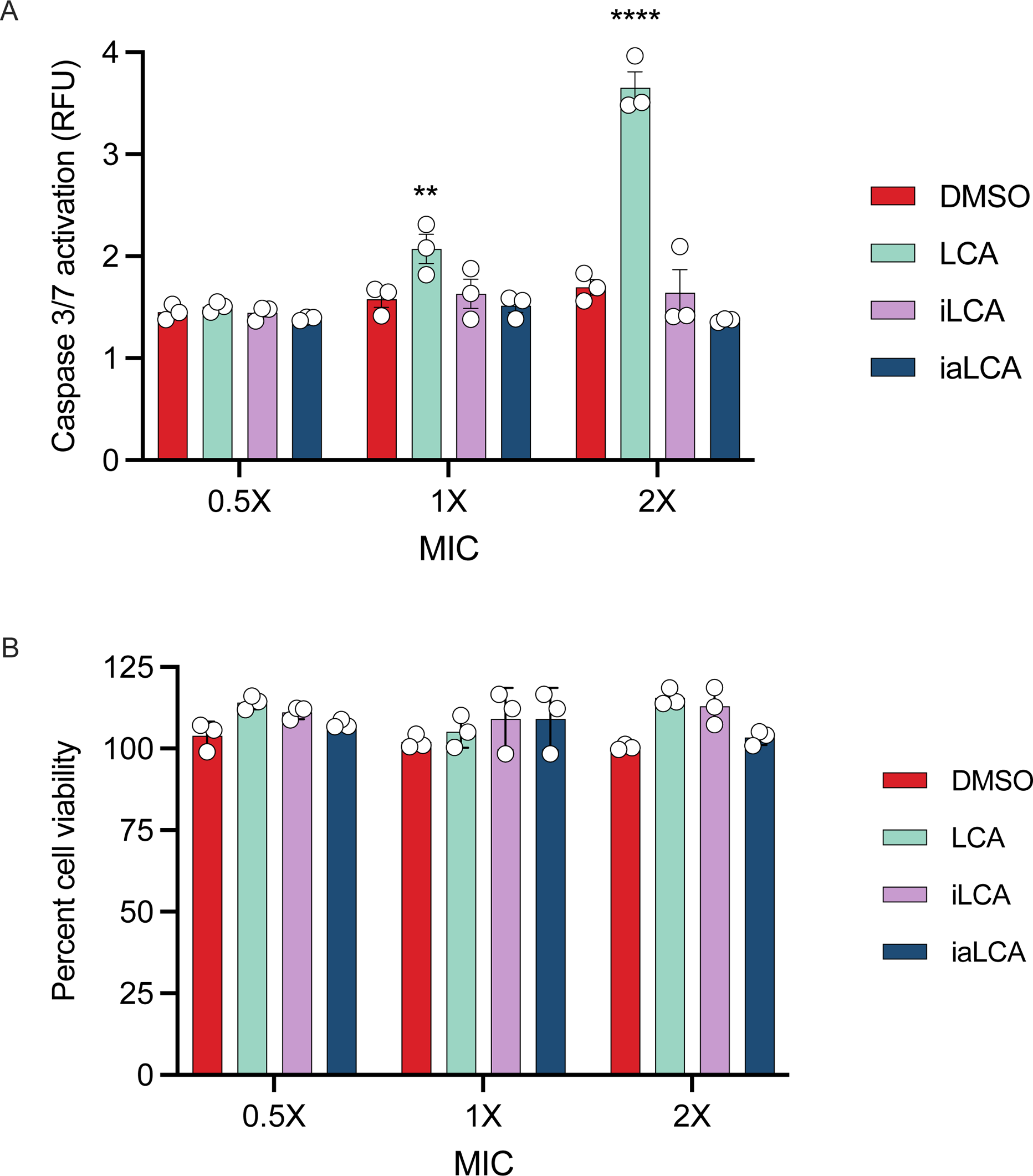
Host cell viability is not affected by LCA derivatives. A modified Caco-2 cell viability assay was used to assess the effects of LCA, iLCA, and iaLCA on host cells. Statistical significance was determined by two-way ANOVA (**, p < 0.01; ****, p < 0.0001). Concentrations used are as follow: LCA (0.01, 0.2, 0.4 mM), iLCA (0.025, 0.05, 0.1 mM), iaLCA (0.01, 0.02, 0.04 mM). Caspase 3/7 activation data (A) represents mean ± SEM of triplicate samples. Percent cell viability (B) data represents mean ± SD of triplicate samples.

## Discussion

As *C. difficile* infections and antibiotic resistance continue to increase, the urgency to develop new antibiotics and or therapeutics that can target the pathogen and spare the surrounding gut microbiota increases. In this study, we examined how the secondary bile acid LCA and its epimers iLCA and iaLCA inhibit different stages of the *C. difficile* lifecycle. Secondary bile acids such as LCA are able to inhibit *C. difficile*, and their loss in the gut is associated with increased susceptibility to CDI [10, 26]. LCA is epimerized by gut bacteria that encode 3⍺– or 3β-HSDH to produce iLCA and iaLCA [15, 24]. As LCA is further biotransformed to its epimers iLCA and iaLCA, the MIC against *C. difficile* is further reduced (Fig. 1, Table 1). The MIC that we observed for LCA, iLCA, and iaLCA have been reported against *C. difficile* before [14, 24, 26, 39]. iaLCA inhibits a variety of Gram-positive bacteria such as *C. difficile* 630, while iLCA was shown to alter the growth kinetics of seven clinically relevant strains of *C. difficile:* R20291, CD196, M86, CF5, 630, BI9, and M120 [24, 26]. Our findings align with this, but indicate that LCA, iLCA, and iaLCA all differentially impact *C. difficile* growth kinetics. Secondary bile acids such as LCA also have detergent like activity that is associated with membrane damage to many gut bacteria [41]. In comparison, iso-epimers of secondary bile acids have less detergent activity, owing to differences in structure and chemistry [31]. This was consistent in our study as LCA and iaLCA were able to disrupt the *C. difficile* membrane, while iLCA did not.

We observed that epimers of LCA significantly affect toxin expression (*tcdA*) in *C. difficile* in a dose dependent manner, while only LCA significantly affects toxin activity. This could be due to LCA’s ability to bind to *C. difficile* toxin TcdB, which induces conformational changes that limit toxin binding and activity [27]. This is also consistent with recent work showing that secondary bile acid extracts from mouse and human stool samples were protective against TcdB-mediated cell damage [43]. These observed differential effects between LCA, iLCA, and iaLCA indicate that these epimers may be acting on different aspects of the *C. difficile* lifecycle, with LCA targeting the activity of the toxin while iLCA and iaLCA prevent the production of toxins via decreased *tcdA* expression. It is not known if the LCA epimers are able to bind to TcdB, and it is also not known how these epimers affect expression of *tcdB*. Future work looking at how these epimers affect *C. difficile* toxin and other virulence factors will be of interest.

The concentrations of bile acids tested in our study were below physiological concentrations, although there has been a wide range observed. LCA has been detected in the intestinal tract at concentrations ranging from 0.001 mM to 10.4 +/− 7.5 mM [29, 30]. Likewise, iLCA has been detected at concentrations ranging from 0 mM to 0.360 mM to 1.0155 +/− 1.1276 mM while iaLCA has been measured at 0.0359 +/− −.1196 mM [29, 30]. It is important to note that the donor sources and types of samples vary greatly between studies. Samples from cecal content or stool were used to quantify and classify bile acids in cited studies, and in conjunction with donor source (live versus deceased), may account for the variability observed [29, 30]. Regardless of donor source, the concentrations of LCA, iLCA, and iaLCA used in this study all fall within the ranges of bile acid concentrations observed in humans.

While secondary bile acid epimers are known to affect *C. difficile* growth and virulence factors, they also play a role in modulating the composition of the indigenous gut microbiota. The presence of iaLCA influences the growth of Gram-positive commensals, including several species of *Clostridia* like *Clostridium sporogenes, Clostridium hiranonis,* and *Clostridium indolis*, however it is less inhibitory toward these commensals than it is to *C. difficile* [24]. Additionally, iaLCA has favors the growth of Gram-negatives such as *Bacteroides,* while impeding the growth of Gram-positives such as *Streptococcus* and *Bifidobacterium* [24]. Likewise, the production of isoDCA (iDCA) favors the growth of *Bacteroides* [31]. Our findings echo this and show that LCA and its epimers are more inhibitory towards *Clostridia* and spare other members of the gut microbiota including the Gram-negatives *Escherichia coli, Bacteroides thetaiotaomicron,* and *Muribaculum intestinale* DSM 28989 (Table 2). It has been hypothesized that the lower toxicity to Gram-negative commensals is due to the lower detergent properties of iDCA when compared to DCA [31]. This reduction in toxicity towards certain members of the gut microbiota has also been hypothesized to be beneficial to the bacteria that produce secondary bile acids and their epimers, such as *Ruminococcus gnavus* or *Eggerthella lenta* [31]. As a whole, LCA, iLCA, and iaLCA produce conditions favorable for the growth of some bile acid modifying commensals while limiting the outgrowth of other gut bacteria.

In addition to modulating the composition of the gut microbiota, epimers of secondary bile acids also play a regulatory role within the host. Bile acids act in several signaling pathways associated with host-bile acid production, immunity, and inflammation. Secondary acids such as LCA bind the farnesoid x receptor (FXR) as ligands and alter the transcription of genes associated with bile acid production and transportation [44, 45]. This causes shifts in the composition and enterohepatic circulation of the bile acid pool. Particularly, the activation of FXR by secondary bile acids has been shown to decrease synthesis of bile acids, which can functionally limit the effects of bile acid toxicity while also shifting the microbiota structure [45–47]. As we observed that LCA can be damaging to host cells below physiological concentrations, it is beneficial to have a level of regulation that can reach secondary bile acids to prevent adverse events [30] (Fig. 6).

Bile acids also can provide protective effects to the host by interacting with immune cells, thereby modulating the host’s immune state. The LCA derivative 3-oxoLCA inhibits differentiation of T_H_17 cells, while iaLCA promotes regulatory T cell (T_reg_) differentiation [48]. Both LCA derivatives limit inflammation in the host through their respective interactions with adaptive immune cells [48]. Furthermore, secondary bile acid activation of the G-protein-coupled receptor TGR5 has been shown to limit inflammation [46]. Patients experiencing gastrointestinal inflammation often have reduced levels of secondary bile acids, their epimers, and the bacteria that are responsible for their production [46]. Loss of commensals that produce iaLCA has been observed in both Crohn’s disease and ulcerative colitis patient cohorts, and a reduction in secondary bile acids present has also been documented in Crohn’s disease patients [30, 33]. The recovery of secondary bile acids and their epimers through FMT, which provides the bacteria that can modify bile acids, has been observed [30]. This suggests that the presence of secondary bile acids, and more importantly their epimers, may be essential for reducing inflammation and proper immunomodulation.

The LCA derivatives iLCA and iaLCA are attractive candidates for future study as therapeutics for CDI and other gastrointestinal disease. Although they can eliminate *C. difficile in vitro* and *in vivo*, there are several challenges that must be addressed before they can be used in humans [24]. The most pressing issue facing the use of secondary bile acids in their epimers is the delivery mechanism. As bile acids travel from the ileum to the colon, they are inherently modified by the indigenous gut microbiota and absorbed, which may alter their efficacy at the target site. Because of this, it may be difficult to provide an accurate dosage to a specific section of the intestinal tract. One potential route to overcome this issue is using the bacteria that make LCA and its epimers. Several groups have identified bacteria that produce iLCA and iaLCA, and this knowledge could be used to rationally design probiotics that allow for the natural production of a desired bile acid [14, 30]. An alternative method of bile acid delivery could be via a genetically engineered strain that secretes a specific bile acid [49]. Future work on LCA and its epimers should determine the optimal dose and delivery mechanism of these bile acids to maximize their therapeutic impact.

In the search for a novel therapeutic that targets *C. difficile*, bile acids have become a viable solution. Epimers of bile acids are particularly attractive as they may provide protection against *C. difficile* while leaving the indigenous gut microbiota largely unaltered. This study shows that iLCA and iaLCA specifically are potent inhibitors of *C. difficile*, affecting key virulence factors including growth, toxin expression and activity. As we move toward the use of bile acids as therapeutics, further work will be required to determine how best to deliver these bile acids to a target site within the host intestinal tract.

## Data availability

Data is available upon request.

## Acknowledgements

SCK was funded by the NCSU Molecular Biology Training Program grant T32 GM008776 through the NIH. CMT was funded by the National Institute of General Medical Sciences of the National Institutes of Health under award number R35GM119438. CMT is a consultant for Ferring Pharmaceuticals and on the Scientific Advisory Board for Ancillia Biosciences.

